# MAK and CCRK kinases regulate kinesin-2 motility in *C. elegans* neuronal cilia

**DOI:** 10.1101/209221

**Authors:** Peishan Yi, Chao Xie, Guangshuo Ou

**Affiliations:** Tsinghua-Peking Center for Life Sciences, School of Life Sciences and MOE Key Laboratory for Protein Science, Tsinghua University, Beijing 100084, China

## Abstract

Kinesin-2 motors power the anterograde intraflagellar transport (IFT), a highly ordered process that assembles and maintains cilia. It remains elusive how kinesin-2 motors are regulated *in vivo*. Here we perform forward genetic screen to isolate suppressors that rescue the ciliary defects in the constitutive active mutation of OSM-3-kinesin (G444E) in *C. elegans* sensory neurons. We identify the *C. elegans* DYF-5 and DYF-18, which encode the homologs of mammalian male germ cell-associated kinase (MAK) and cell cycle-related kinase (CCRK). Using time-lapse fluorescence microscopy, we show that DYF-5 and DYF-18 are IFT cargo molecules and are enriched at the distal segments of sensory cilia. Mutations of *dyf-5* and *dyf-18* generate the elongated cilia and ectopic localization of kinesin-II at the ciliary distal segments. Genetic analyses reveal that *dyf-5* and *dyf-18* are also important for stabilizing the interaction between IFT particle and OSM-3-kinesin. Our data suggest that DYF-5 and DYF-18 act in the same pathway to promote handover between kinesin-II and OSM-3 in sensory cilia.

## Introduction

Cilia are microtubule-based organelles that play important roles in signal sensation and motility (Ishikawa and Marshall, 2017). The assembly and maintenance of cilia rely on the intraflagellar transport (IFT), a process first discovered in *Chlamydomonas* and later found to exist in various ciliated organisms (Ishikawa and Marshall, 2011; Kozminski et al., 1993; Rosenbaum and Witman, 2002; Scholey, 2003; Taschner and Lorentzen, 2016). This process exploits multiprotein complexes, IFT particles, to transport various ciliary components bidirectionally between the ciliary base and tip (Ishikawa and Marshall, 2011; Taschner and Lorentzen, 2016). The anterograde IFT, namely the transport of IFT components from the ciliary base to the tip is driven by the kinesin-2 motors, whereas the retrograde transport of IFT components from the tip to the base is powered by the cytoplasmic dynein-2.

The nematode *Caenorhabditis elegans* is a valuable model organism in identifying and investigating IFT components including kinesin-2 motors. The heterotrimeric kinesin-II which is composed of two different motor subunits and a kinesin-associated subunit (KAP) is employed by most species to power the anterograde IFT (Scholey, 2013). The assembly of *C. elegans* sensory cilia also requires another homodimeric kinesin-2 OSM-3 (Evans et al., 2006; Scholey, 2013; Signor et al., 1999; Snow et al., 2004). In *C. elegans* amphid and phasmid sensilla, cilia are assembled into bipartite structures that contain a middle segment with nine doublet microtubules and a distal segment with nine singlet microtubules (Inglis et al., 2007). Kinesin-II is restricted to the middle segments, while OSM-3 moves along the entire cilia (Ou et al., 2005; Prevo et al., 2015; Snow et al., 2004). Genetic studies demonstrate that OSM-3 and kinesin-II cooperate to build up the middle segments whereas OSM-3 alone assembles the distal segments of sensory cilia (Ou et al., 2005; Snow et al., 2004). The handover mechanism of kinesin-II and OSM-3 in anterograde transport has been recently demonstrated : kinesin-II is required for efficient import of IFT particles from cell body into the ciliary compartment whereas OSM-3 mediates the long distance transport along the whole cilia (Prevo et al., 2015). However, how this process is regulated and how kinesin-2motors are controlled to move bidirectionally along the axoneme are not well understood.

The precise localization and regulation of kinesin motor proteins are essential for the coherent intracellular transport (Scholey, 2013; Verhey and Hammond, 2009). The activity of kineins-2 motors can be modulated through an autoinhibitory mechanism. For instance, OSM-3 and its mammalian homolog KIF17 are autoinhibited by residing in a compact conformation with involvement of interactions between the tail and motor domains (Hammond et al., 2010; Imanishi et al., 2006). Interfering these interactions or mimicking cargo loading, which is supposed to maintain the extended active conformation, produces enhanced motility and processivity. It is unclear how the autoinhibitory mechanism applies to kinesin-2 activity during ciliogenesis. A simple model would be that OSM-3 become active once it is docked onto IFT particles. Indeed, loss of IFT component DYF-1/IFT70 inactivates OSM-3 motility (Ou et al., 2005). However, no direct interaction between DYF-1 and OSM-3 has been reported and DYF-1 fails to activate OSM-3 *in vitro* (Imanishi et al., 2006). Another supporting evidence comes from the analyses of the *sa125* mutation of OSM-3 which converts a Glycine to Glutamate (G444E) at the hinge region and relieves the autoinhibited state without cargo binding in single-molecule assays (Imanishi et al., 2006). Interestingly, *sa125* mutant worm forms defective cilia as *osm-3* null alleles (Snow et al., 2004). OSM-3(G444E) may become constitutively active and fail to bind IFT cargos, making it unable to be assembled or travel into cilia (Imanishi et al., 2006).

The autoinhibitory mechanism of the classical heterotrimeric kinesin-II is less understood, although its folded and extended conformations are observed under electron microscopy. An *in vitro* study showed that mutations in the coiled coil region increase the motility and processitivity of heterodimerized motor subunits of kinesin-II (Brunnbauer et al., 2010; Wedaman et al., 1996). Phosphorylation can provide another layer of regulation of kinesin-II activity and localization in cilia. The *Chlamydomonas* kinesin-II motor subunit FLA8/KIF3B is phosphorylated and inactivated by the calcium-dependent kinase CDPK1, which inhibits IFT entry probably by disrupting the interactions between kinesin-II and IFT particles (Liang et al., 2014). Other kinases including the cell cycle-related kinase (CCRK), the ros-cross hybridizing kinase (RCK) family members (MAK, ICK and MOK) and their homologs in *Chlamydomonas*, *C. elegans* and other species are implicated in the inhibition of kinesin-II due to the consistent long-cilia phenotypes and compromised IFT in their absence (Asleson and Lefebvre, 1998; Bengs et al., 2005; Berman et al., 2003; Broekhuis et al., 2014; Burghoorn et al., 2007; Chaya et al., 2014; Moon et al., 2014; Omori et al., 2010; Tam et al., 2007; Yang et al., 2013). MAK and ICK are substrates of CCRK and ICK is reported to phosphorylate mammalian kinesin-II motor subunit KIF3A (Chaya et al., 2014; Fu et al., 2006; Wang and Kung, 2012; Yang et al., 2013). However, CCRK and its homologs do not localize to cilia (Ko et al., 2010; Phirke et al., 2011; Tam et al., 2007; Yang et al., 2013), which is inconsistent of the site of their action. MAK/ICK and their homologs are reported to localize to the whole cilia, at ciliary base or at ciliary tip; the localization of ICK can be either dependent or independent of its kinase activity (Berman et al., 2003; Broekhuis et al., 2014; Burghoorn et al., 2007; Chaya et al., 2014; Moon et al., 2014; Omori et al., 2010).

Here we first report that the OSM-3 hinge mutation G444E is insufficient to constitutively activate it in cilia. In contrast, OSM-3(G444E) is passively transported by kinesin-II likely together with IFT particles. In our attempt to isolate suppressors of the ciliary defects in *osm-3(G444E)*, we have identified two kinases DYF-5/MAK and DYF-18/CCRK. We show that DYF-5 and DYF-18 are IFT cargo molecules that are enriched at the distal ciliary segment, which in turn restrict the kinesin-II within the middle segment. Our genetic analyses also suggest that DYF-5 and DYF-18 act in the same pathway to regulate the localization of kinesin-II and coordination between kinesin-II and OSM-3. We propose a model that DYF-5 and DYF-18 promote the handover between kinesin-II and OSM-3 and turnaround or docking of IFT particles likely through remodeling of IFT machineries during retrograde IFT at the distal segment.

## Results

### OSM-3(G444E) is passively transported by kinesin-II

The autoinhibition model infers that the G444E mutation at the hinge region (H2) relieves autoinhibition of OSM-3 without cargo loading (Imanishi et al., 2006) (Figure 1A). To test the possibility *in vivo*, we overexpressed GFP-labeled OSM-3(G444E) under the control of a ciliated neuron-specific promoter (*Pdyf-1*) in the wild type (WT) *C. elegans* (Figure 1B-D). Surprisingly, we found that both amphid and phasmid sensory cilia are severely truncated and lose the distal segments in the transgenic animals, phenocopying the G444E mutant *osm-3(sa125)* or the null mutant *osm-3(p802)*. Overexpression of OSM-3 with a small deletion at the hinge region (ΔH2) also resulted in the same phenotypes (Figure 1E). In the remaining middle segments, OSM-3(G444E)::GFP moved at 0.63 ± 0.16 μm/s (mean ± s.d., N=184) in the anterograde direction, which is smaller than the WT velocity (~0.7 μm/s at the distal segments and ~1.3 μm/s at the distal segments) (Snow et al., 2004). The mCherry labeled IFT particle-B subcomplex component OSM-6/IFT52 travelled at a similar speed (0.65. ± 0.10 μm/s, mean ± s.d., N=123). These results indicate that OSM-3(G444E) is not constitutively active but may be still associated with the IFT particles and that the mutant protein could interfere the function of the endogenous WT OSM-3.

**Figure 1.**
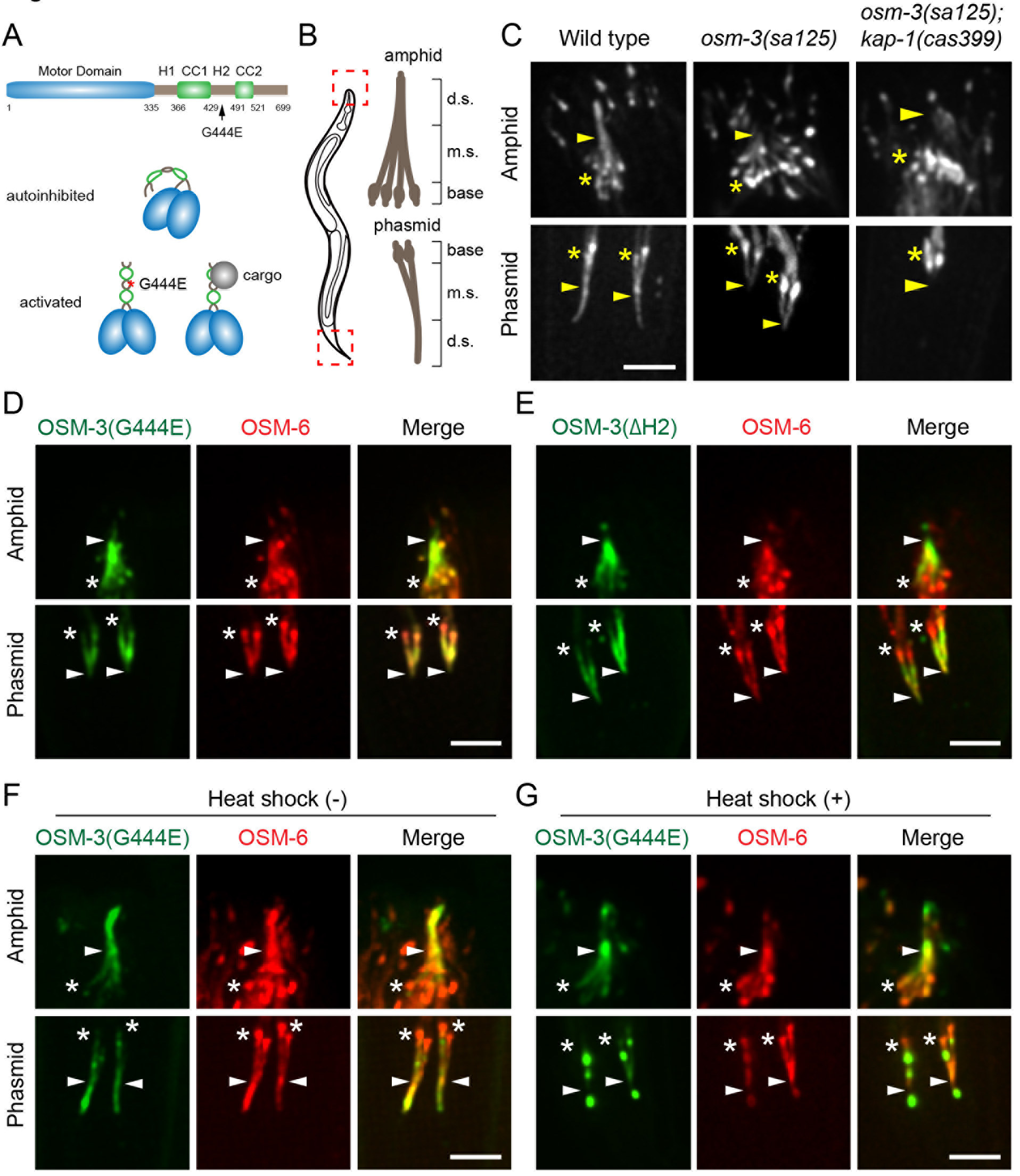
OSM-3(G444E) impairs cilium formation. (A) Domain structures of OSM-3 and the autoinhibition model. The motor domain (blue), coiled-coils (CC1 and CC2) and hinge regions (H1 and H2) are indicated. At the autoinhibited state, OSM-3 tail folds back to generate a compact conformation. Cargo loading or the G444E mutation at hinge 2 relieves autoinhibition. (B) Schematic depiction of the *C. elegans* amphid and phasmid cilia. Ten or two sensory cilia are aligned into the amphid or phasmid channels (dashed boxes and enlarged on the right), respectively. Each cilium contains a ciliary base, a middle segment (m.s.) and a distal segment (d.s.). The distal segments are exposed to the environment (not shown). (C) Amphid (up) and phasmid (below) cilia in the wild type, G444E mutant *osm-3(sa125)* and *osm-3; kap-1* double mutant animals. Cilia are marked with GFP-tagged IFT52/OSM-6. Star, ciliary base; arrowhead, junction between the middle and distal segments; scale bar, 5 μm (the same in Figure 1D-G). (D) Ciliary phenotypes in OSM-3(G444E) transgenic animals. OSM-3(G444E)::GFP and OSM-6::mCherry are expressed under the control of a ciliated neuron-specific promoter *Pdyf-1*. (E) Ciliary phenotypes in OSM-3(ΔH2) transgenic animals. OSM-3(ΔH2)::GFP and OSM-6::mCherry are expressed under the control of *Pdyf-1*. ΔH2, deletion of the hinge 2. (F) Ciliary phenotypes in OSM-3(G444E) transgenic animals. OSM-3(G444E)::GFP is expressed under the control of a heat shock promoter *Phsp-16.2* without heat shock treatment. OSM-6::mCherry is expressed under the control of *Pdyf-1* promoter. (G) Ciliary phenotypes in OSM-3(G444E) transgenic animals. OSM-3(G444E)::GFP is expressed under the control of *Phsp-16.2* after a one-hour heat shock treatment at 33 °C.

To further test this possibility, we expressed OSM-3(G444E)::GFP under the control of a heat-shock-inducible promoter (P*hsp-16.2*). Low level expression of OSM-3(G444E)::GFP was induced at room temperature without heat shock, which did not affect cilium assembly, allowing the measurement of velocity in the intact cilia (Figure 1F). In the middle and distal segments, OSM-3(G444E)::GFP moved at 0.68 ± 0.15 μm/s (mean ± s.d., N=144) and 1.03 ± 0.17 μm/s (mean ± s.d., N=116), respectively, which are slower than the WT motor. Robust expression of OSM-3(G444E)::GFP induced by heat shock treatment caused severe truncation of the distal segments as caused by expressing OSM-3(G444E)::GFP under the control of *Pdyf-1* (Figure 1E & 1G). Furthermore, OSM-3(G444E)::GFP formed aggregates at the ciliary base and tip (Figure 1G). These results indicate that OSM-3(G444E) is not fully active and can impair the activity of endogenous OSM-3. The dominant negative effects may be caused by the formation of heterodimers by the WT and mutant protein or competition for cargo binding.

To exclude the effects of endogenous OSM-3 on the measurement of the speed of OSM-3(G444E), we expressed OSM-3(G444E)::GFP and mCherry labeled kinesin-II subunit KAP-1 in the G444E mutant *osm-3(sa125)*. OSM-3(G444E)::GFP and KAP-1::mCherry move at 0.56 ± 0.12 μm/s (mean ± s.d., N=138) and 0.57 ± 0.11 μm/s (mean ± s.d., N=62), respectively, suggesting that they may move together. Because OSM-3(G444E) alone is unable to assemble cilia in the absence of kinesin-II as evidenced by completely loss of cilia in *kap-1(null); osm-3(sa125)* double mutant (Figure 1C) and kinesin-II and OSM-3 are not directly associated (Ou et al., 2005), OSM-3(G444E) may be inactive and transported by kinesin-II together with IFT particles.

### Mutations of *dyf-5* and *dyf-18* suppress ciliary defects in *osm-3(sa125)*

*In vitro* results and our data suggest that G444E may affect the conformational state of OSM-3 (Imanishi et al., 2006). We speculate that G444E disrupts essential inter-or intra-molecular interactions, which may be rescued by other mutations of OSM-3 or its regulators. To seek for such mutations, we performed a large scale genetic screen to identify mutations that can suppress the ciliary defects in the G444E mutant *osm-3(sa125)* background (Figure 2A). We identified two genes *dyf-5* and *dyf-18*, whose mutations rescued the dye filling defects (Dyf) of *osm-3(sa125)* (Figure 2B & 2C). *dyf-5* encodes the homolog of mammalian RCK members and *Chlamydomonas* long-flagella 4 (LF4) and *dyf-18* is homologous to the mammalian CCRK and *Chlamydomonas* long-flagella 2 (LF2). We obtained three alleles of *dyf-5* and eight alleles of *dyf-18* (Figure 2B & 2C). All mutations introduce amino acid changes in the conserved kinase domains except two alleles of *dyf-18 cas387* and *cas392* (Figure S1A). *cas387* is a G-to-A conversion at the splicing site following exon five and *cas392* converts a conserved cytosine to thymine in the X-box motif of 5’UTR. The X-box motif was previously demonstrated to be essential for ciliary gene expression (Efimenko et al., 2005).

**Figure 2.**
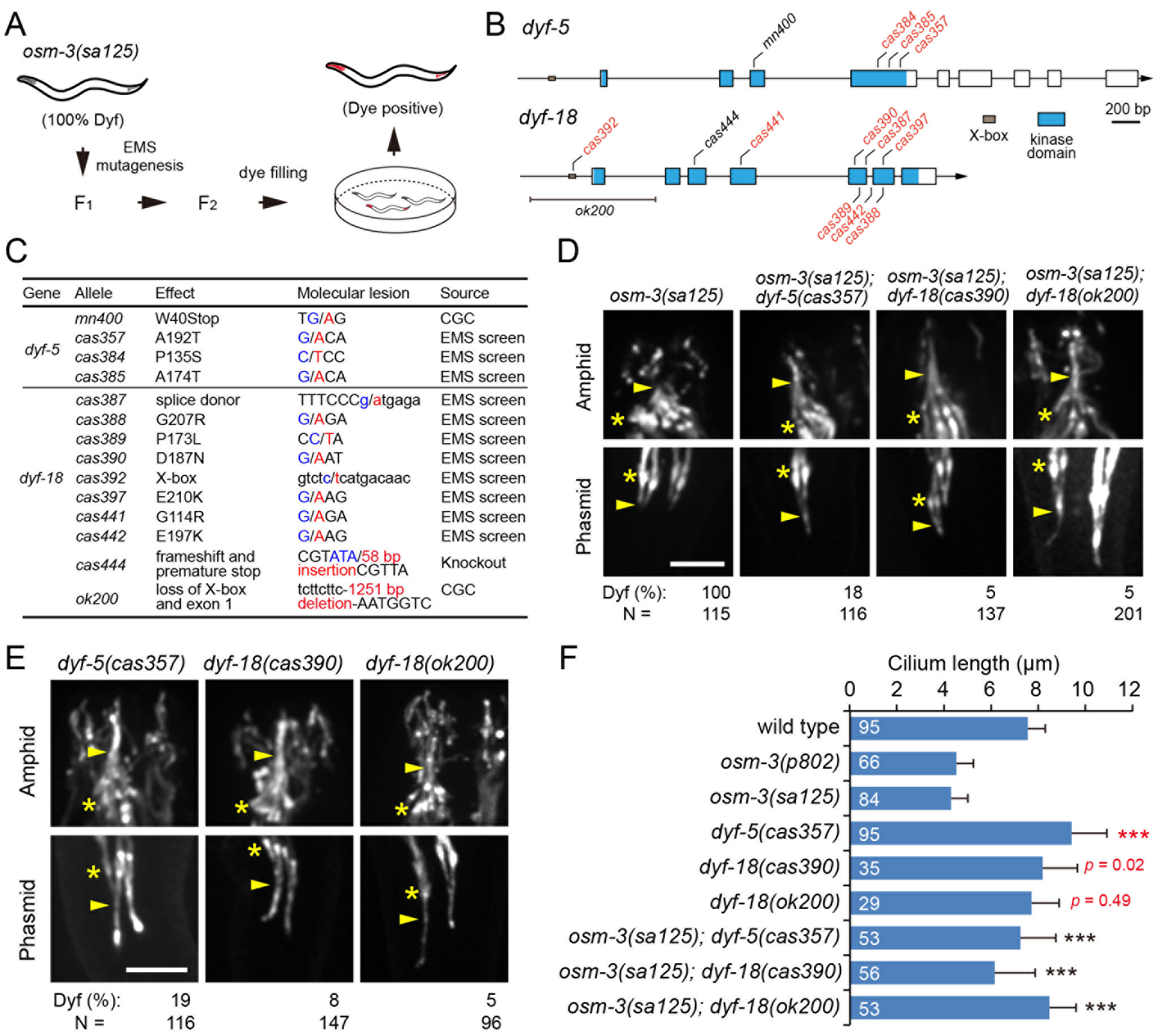
Mutations of *dyf-5* and *dyf-18* suppress ciliary defects in *osm-3(sa125)* mutant. (A) Flowchart of the suppressor screening. *osm-3(sa125)* animals are 100% dye filling defective (Dyf). Animals at late L4 stage were treated with ethyl methanesulfonate (EMS). F2 progenies were screened by filling fluorescent dye DiI. Dye positive animals were considered as the putative suppressors. The ciliary phenotypes were verified under microscopy. Mutations were subjected to genetic mapping (see Materials and Methods). (B) Genomic structures of *dyf-5* (isoform a) and *dyf-18*. Exons are marked with large boxes. Blue regions indicate the coding sequence of kinase domain. Small boxes in grey are X-box motifs within the promoters. Alleles isolated by EMS screening are in red. Other alleles are in black. Scale, 200 bp. (C) Details of *dyf-5* and *dyf-18* alleles. *cas444* was constructed by CRISPR-Cas9-based genome editing. CGC, *Caenorhabditis* Genetics Center. (D) Ciliary phenotypes in *osm-3* single mutant, *osm-3; dyf-5* and *osm-3; dyf-18* double mutants. Cilia are visualized with GFP-tagged IFT52/OSM-6. Dyf, dye filling defective; Star, ciliary base; arrowhead, junction between the middle and distal segments; scale bar, 5 μm (the same in Figure 2E). (E) Ciliary phenotypes in *dyf-5* and *dyf-18* single mutants. (F) Quantification of cilium length (mean ± s.d.). Numbers of worms are indicated in the bars. Red stars or *p* values represent comparisons between the wild type and mutant animals; black stars represent comparisons between the *osm-3(sa125)* single mutant and the double mutants. ***, *p* < 0.001, two-tailed Student’s *t* test.

We next examined the ciliary phenotypes in these suppressors. Using GFP labeled OSM-6/IFT52, a component of IFT-particle B subcomplex, we found that the distal segments in the amphid and phasmid cilia are restored and cilium length increased from 4.3 μm up to ~7.3 μm (Figure 2D & 2F). Consistent phenotypes were also observed when using two ciliated-neuron-specific tubulin markers (Figure S2). By introducing two other independent alleles *ok200* (a deletion allele from *Caenorhabditis* Genetics Center) or *cas444* (a small deletion and insertion allele constructed by CRISPR-Cas9 knockout) of *dyf-18* into *osm-3(sa125)* mutant, we found the similar phenotypes, demonstrating that *dyf-18* is a true suppressor gene (Figure 2D & 2F). As overexpression of WT *dyf-5* causes Dyf phenotype (Burghoorn et al., 2007), we cannot perform a rescue experiment to demonstrate *dyf-5* is responsible for the suppression. However, the isolation of multiple independent alleles suggests that *dyf-5* can be a suppressor. In addition, *dyf-5(cas357)* decreased the Dyf phenotype of *osm-3(p802)* null mutant from 100% to 43% (N=141). Interestingly, the null mutation *mn400* does not suppress either *osm-3(sa125)* or *osm-3(p802)*, suggesting that DYF-5 may have additional functions.

Next, we examined the ciliary morphology in the *dyf-5* and *dyf-18* single mutants. We found the cilium length was remarkably increased in the *dyf-5* single mutant and to a lesser extend in *dyf-18* mutants, which is consistent with the long-cilia phenotypes observed in the previous studies (Burghoorn et al., 2007; Phirke et al., 2011) or in the absence of their counterparts in *Chlamydomonas* and mammalian cells (Asleson and Lefebvre, 1998; Berman et al., 2003; Broekhuis et al., 2014; Moon et al., 2014; Omori et al., 2010; Tam et al., 2007; Yang et al., 2013) (Figure 2E & 2F). However, we only observed subtle Dyf phenotypes in each mutants, indicating that these mutations do not completely block IFT and ciliogenesis (Figure 2E). Indeed, only mild accumulation of IFT particles at the ciliary tip were observed and IFT was detectable along the full length cilia, although the speed is much slower than that in WT animals (Figure 2E, Table 1). Given that *dyf-5* null mutant has strong ciliary defects, we speculate that these *dyf-5* mutations cause weak loss-of-function.

**Table 1.** Summary of IFT velocities. Velocities were quantified using GFP labeled IFT52/OSM-6 (mean ± s.d., μm/s). Numbers of particles used for quantification are indicated in brackets. m.s., middle segment; d.s., distal segment. N/A, not available.

| Genotype | Anterograde (m.s.) | Anterograde (d.s.) | Retrograde |
| --- | --- | --- | --- |
| *Wild type* | 0.76 ± 0.12 (302) | 1.29 ± 0.24 (206) | 1.24 ± 0.20 (88) |
| *osm-3(p802)* | 0.58 ± 0.07 (332) | None | N/A |
| *osm-3(sa125)* | 0.61 ± 0.08 (208) | None | N/A |
| *dyf-5(cas357)* | 0.67 ± 0.21 (316) | 0.82 ± 0.20 (208) | 1.19 ± 0.28 (255) |
| *dyf-18(cas390)* | 0.76 ± 0.21 (148) | 0.95 ± 0.25 (128) | 1.25 ± 0.31 (142) |
| *dyf-18(ok200)* | 0.63 ± 0.10 (149) | 0.67 ± 0.08 (74) | 1.24 ± 0.20 (82) |
| *osm-3(sa125); dyf-5(cas357)* | 0.65 ± 0.10 (322) | 0.64 ± 0.14 (77) | 1.20 ± 0.27 (99) |
| *osm-3(sa125); dyf-18(cas390)* | 0.57 ± 0.09 (150) | 0.58 ± 0.09 (66) | 1.14 ± 0.18 (42) |
| *osm-3(sa125); dyf-5(cas357); dyf-18(cas390)* | 0.59 ± 0.09 (183) | undetectable | 1.10 ± 0.24 (53) |
| *kap-1(cas399)* | 1.24 ± 0.19 (174) | 1.26 ± 0.30 (80) | N/A |
| *kap-1(cas399); dyf-18(ok200)* | 0.92 ± 0.20 (130) | 1.27 ± 0.36 (121) | N/A |
| *kap-1(cas399); dyf-5(cas357)* | 1.28 ± 0.22 (181) | 1.31 ± 0.22 (148) | N/A |

### DYF-5 and DYF-18 restrict kinesin-II to the middle segment

Previous study has shown that loss of *dyf-5* led to the ectopic entry of kinesin-II into the distal segments (Burghoorn et al., 2007). We asked whether the suppression by *dyf-5* and *dyf-18* was a consequence of functional compensation by kinesin-II in the distal segments. Since the three alleles of *dyf-5* have the same phenotypes and *ok200* is putatively null, we used *cas357* and *ok200* to investigate the roles of *dyf-5* and *dyf-18* in ciliogenesis, respectively. By examining a kinesin-II marker KAP-1::GFP, we found kinesin-II ectopically entered the distal segments in both mutants (Figure 3A). The same localization pattern was observed when we used the GFP labeled endogenous kinesin-II motor subunit KLP-20 (Figure 3B). To further test whether kinesin-II is required for suppression of *osm-3(sa125)*, we generated the *dyf-5(cas357)*; *kap-1(cas399)*; *osm-3(sa125)* triple mutant animal. As expected, this mutant was 100% Dyf, which indicates that kinesin-II is required for the suppression. Compared with *kap-1(cas399)*; *osm-3(sa125)* double mutant (Figure 1C), the triple mutant formed the middle segments and IFT was detectable at the speed of 0.53 ± 0.09 μm/s (mean ± s.d., N=95), suggesting that *dyf-5(cas357)* partially restores the motility of OSM-3(G444E) (Figure 3C).

**Figure 3.**
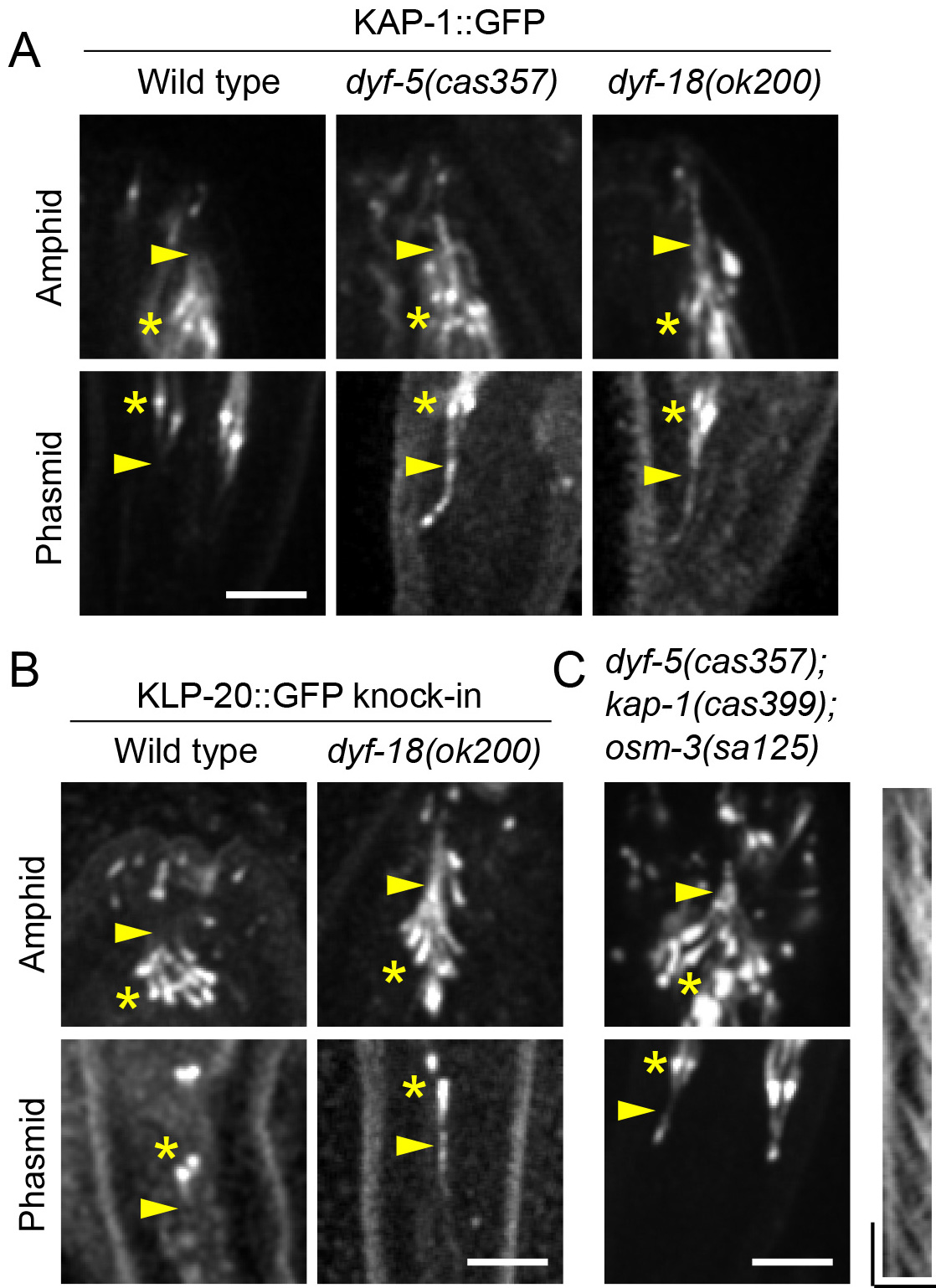
Kinesin-II enters the distal segments in *dyf-5* and *dyf-18* mutants. (A) Localization of GFP labeled kinesin-II accessory subunit (KAP-1) in wild type and mutant worms. Star, ciliary base; arrowhead, junction between the middle and distal segments; scale bar, 5 μm (the same in Figure 3B). (B) Localization of GFP labeled kinesin-II motor subunit (KLP-20) in wild type and *dyf-18* mutant worms. (C) Ciliary phenotypes in *dyf-5; kap-1; osm-3* triple mutant animals. Cilia are marked with GFP-tagged IFT52/OSM-6. Kymograph shows IFT in the middle segments. Star, ciliary base; arrowhead, junction between the middle and distal segments; scale in image, 5 μm; scale in kymographs, horizontal 5 μm; vertical, 5 sec.

### DYF-5 and DYF-18 fine-tune the coordination between kinesin-II and OSM-3

Next, we addressed how DYF-5 and DYF-18 regulate OSM-3-kinesin. In *dyf-5* and *dyf-18* single mutants, the IFT velocities at the middle and distal segments were both reduced, indicative of reduced OSM-3 motility (Table 1). By contrast, the retrograde IFT speed was normal (Table 1). To quantify the velocity of OSM-3-driven IFT alone, we generated the *dyf-5*; *kap-1* and *dyf-18*; *kap-1* double mutants. Unexpectedly, we found that IFT velocities were largely comparable to that in the *kap-1* single mutant where OSM-3 drives IFT at the speed of ~1.3 μm/s (Snow et al., 2004), except a slower speed in the middle segments of the *dyf-18*; *kap-1* double mutant (Figure 4A-4C). These data suggest that DYF-5 and DYF-18 may not affect OSM-3 activity but may be involved in the stabilization of the interaction between OSM-3 and cargo molecules. In support of this notion, we observed a decrease of IFT frequency in *dyf-5*; *kap-1* and *dyf-18; kap-1* double mutants (Figure 4D). We reasoned that in *dyf-5* and *dyf-18* single mutants, the active confirmation of OSM-3 maynot be maintained and that IFT particles were predominantly transported by the slower of kinesin-II. Together with the observation of ectopic kinesin-II localization, we propose that DYF-5 and DYF-18 function to promote the handover of IFT particles between these two motors. Loss of their function results in the failure of undocking of kinesin-II and docking of OSM-3 at the distal segments, which is consistent with previous findings (Burghoorn et al., 2007).

**Figure 4.**
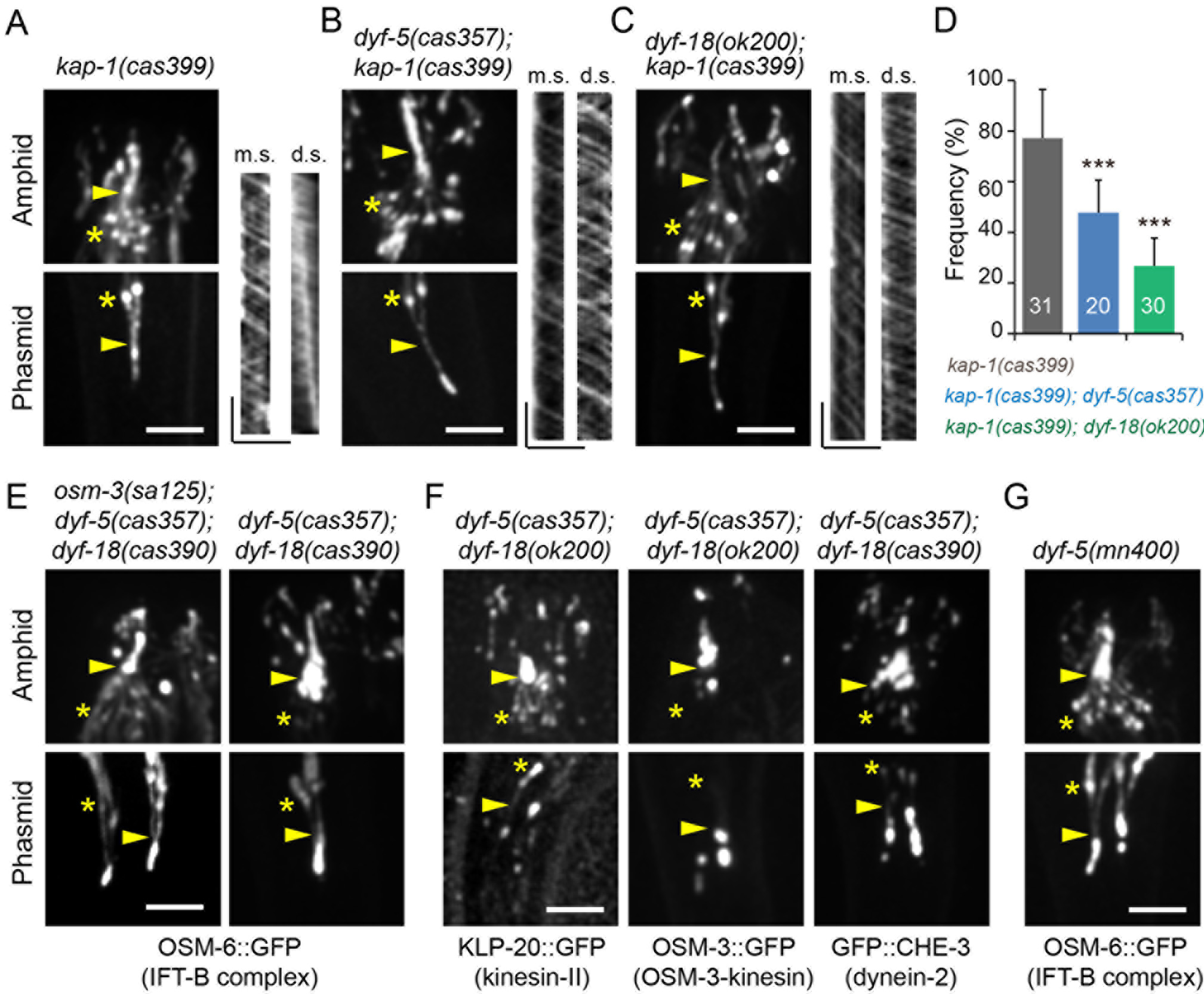
*dyf-5* and *dyf-18* regulate IFT particle-OSM-3 binding and recycling of IFT components. (A) Ciliary phenotypes in kinesin-II’s *kap-1* mutant. Cilia are marked with GFP-tagged IFT52/OSM-6. Kymographs show IFT in the middle (m.s.) and distal (d.s.) segments. Star, ciliary base; arrowhead, junction between the middle and distal segments; scale in image, 5 μm; scale in kymographs, horizontal 5 μm; vertical, 5 sec (the same in Figure 4B & 4C). (B) Ciliary phenotypes in *dyf-5; kap-1* double mutant. (C) Ciliary phenotypes in *dyf-18; kap-1* double mutant. (D) IFT frequencies in *kap-1*, *kap-1; dyf-5* and *kap-1; dyf-18* mutants (mean ± s.d.). Numbers of movies used for quantification are shown in the bars. ***, *p* < 0.001, two-tailed Student’s *t* test. (E) Ciliary phenotypes in *osm-3; dyf-5; dyf-18* and *dyf-5; dyf-18* mutants. Star, ciliary base; arrowhead, junction between the middle and distal segments; scale bar, 5 μm (the same in Figure 4F & 4G). (F) Ciliary phenotypes in *dyf-5; dyf-18* double mutants. (G) Ciliary phenotypes in *dyf-5* null mutant.

### Mutations of *dyf-5* and *dyf-18* impair recycling of IFT components

The similar ciliary phenotypes in *dyf-5* and *dyf-18* mutants promoted us to examine whether they function in the same pathway. We generated the *osm-3(sa125); dyf-5(cas357); dyf-18(cas390)* triple mutant animal. This mutant was completely Dyf and IFT particles were strongly accumulated at the distal segments (Figure 4E), and the cilium length is comparable to that of *osm-3(sa125); dyf-5(cas357)* or *osm-3(sa125); dyf-18(cas390)* double mutant animals. We also observed ectopic localization of kinesin-II in *dyf-5; dyf-18* double mutant (Figure 4F). These data suggest that DYF-5 and DYF-18 act in the same pathway on cilium assembly and regulation of kinesin-II.

We further explored the accumulation of IFT particles at the distal segments of the *dyf-5(cas357); dyf-18(cas390)* double mutants. Only a small proportion of IFT particles appeared to be accumulated in the ciliary tips of each single mutant; however, kinesin-II, OSM-3, dynein-2 and IFT particles all formed remarkable aggregates in the distal segments of *dyf-5; dyf-18* double mutants (Figure 4F). Retrograde velocity is undetectable in the distal segments due to the strong aggregation, but is normal in the middle segments or in the cilia of *dyf-5* or *dyf-18* single mutants (Table 1). These results suggest mutations of *dyf-5* and *dyf-18* affect IFT particle turnaround or docking onto dynein-2 in the distal segments, but not in the middle segments. Considering the similar phenotypes observed in *dyf-5* null mutant and *dyf-5(cas357); dyf-18* double mutants (Figure 4G), we speculate that the enhanced phenotypes in the double mutants may not be resulted from the inactivation of parallel pathways, but represent the different extents of DYF-5 loss of function.

### DYF-5 and DYF-18 are IFT cargo molecules

To determine how DYF-5 and DYF-18 function in cilia, we examined their ciliary localization patterns. We constructed a transgenic animal expressing GFP-tagged DYF-18 that fully rescued the suppression phenotypes in *osm-3(sa125); dyf-18(cas390)* mutant (Figure 5A-B). The GFP fluorescence of DYF-18::GFP distributes along both amphid and phasmid cilia, with enrichment at the distal segments and deprivation from the ciliary base (Figure 5A & 5D). As overexpression of DYF-5 causes strong ciliary defects (Burghoorn et al., 2007), we generated GFP knock-in at the carboxyl terminus of endogenous isoform a (DYF-5a), which does not show any apparent ciliary abnormality. Although the endogenous expression level of DYF-5a::GFP is weak in cilia, we detected an enrichment of the GFP fluorescence in the distal segments and ciliary tips (Figure 5C & 5D). To exclude the effects of kinase activity on cilium formation, we generated transgenic animals expressing GFP-tagged kinase-dead DYF-5(K40A) or DYF-18(K37A). Both proteins are localized in cilia and show an enrichment at the distal segments, except for a lesser extent of enrichment compared to the WT proteins (Figure 5E & 5F). Importantly, our kymograph analyses detected the bidirectional IFT of each protein, indicating that they are transported by IFT. Previous studies reported that DYF-5 and DYF-18 localize to the ciliary base (Burghoorn et al., 2007; Phirke et al., 2011), which may be resulted from overexpression or different experimental conditions. Our usage of genome-editing technique and the ciliary phenotype support the action sites of DYF-5 and DYF-18 at the distal segments or ciliary tips.

**Figure 5.**
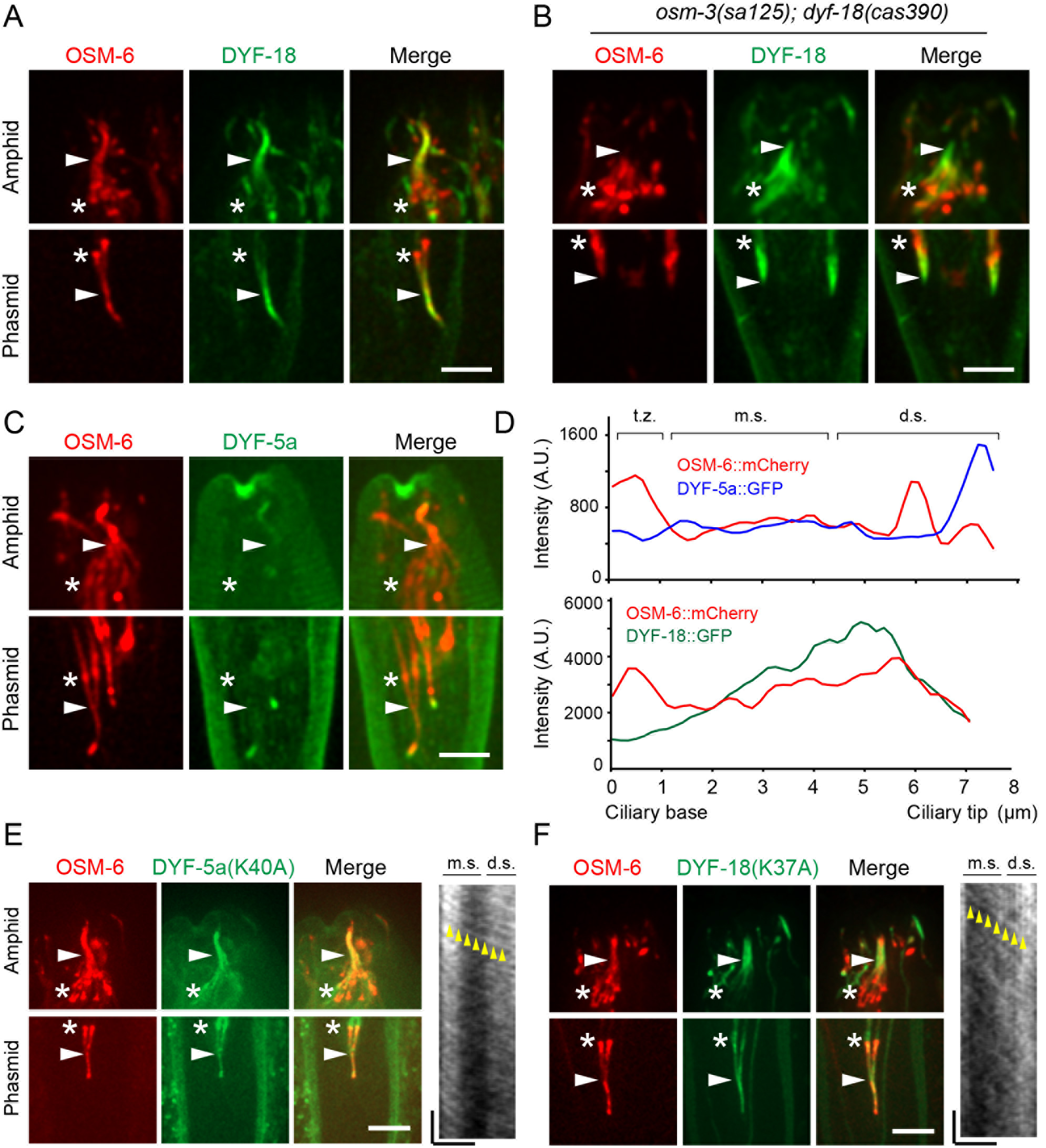
DYF-5 and DYF-18 localize to cilia and undergo IFT. (A) Localization of DYF-18 in cilia. DYF-18::GFP and OSM-6::mCherry are expressed under the control of the *Pdyf-1* promoter. Star, ciliary base; arrowhead, junction between the middle and distal segments; scale bar, 5 m (the same in Figure 5B & 5C). (B) Expression of DYF-18::GFP rescued the *dyf-18*-dependent suppression of *osm-3(sa125)*. OSM-6::mCherry marks cilia. (C) Endogenous localization of DYF-5a::GFP in the cilia of DYF-5a::GFP knock-in animals. (D) Representative fluorescence intensity of DYF-5a::GFP and DYF-18::GFP along cilia. t.z., transition zone; m.s., middle segment; d.s., distal segment. A.U., arbitrary unit. (E) Ciliary localization and IFT of DYF-5a::GFP bearing the kinase-dead mutation (K40A). DYF-5a(K40A)::GFP and OSM-6::mCherry are expressed under the control of the *Pdyf-1* promoter. Star, ciliary base; white arrowhead, junction between the middle and distal segments; yellow arrowheads indicate a representative movement; scale in image, 5 μm; scale in kymographs, horizontal 5 μm; vertical, 5 sec. m.s., middle segment; d.s., distal segment (the same in Figure 5F). (F) Ciliary localization and IFT of DYF-18::GFP bearing the kinase-dead mutation (K37A). DYF-18(K37A)::GFP and OSM-6::mCherry are expressed under the control of the *Pdyf-1* promoter.

### Kinase activity and ciliary localization are required for DYF-18 function

The isolation of multiple mutations affecting conserved residues in the kinase domains of DYF-5 and DYF-18 underscores the dependence of kinase activity in their ciliary functions. To further test this possibility, we used the kinase-dead constructs DYF-5(K40A) and DYF-18(K37A) to rescue *osm-3; dyf-5* or *osm-3; dyf-18* mutants, respectively. As expected, both constructs failed to rescue the ciliary phenotypes (Figure 6A & 6B). Considering that DYF-18 is excluded from the ciliary base, we asked whether the accurate ciliary localization pattern is important for DYF-18 function. To this end, we fused DYF-18 to the transitionzone protein MKSR-2. This chimeric protein was remarkably localized at the ciliary base and transition zone with only dim signal in the cilia (Figure 6C). We found 11 out of 14 worms showed the alignment defect of phasmid cilia, which has never been observed in the WT or DYF-18 overexpression lines (Figure 6C). These data indicate the importance of the proper ciliary localization of DYF-18 in cilium formation. This phenotype was also detectable in the *dyf-5* null mutant, suggesting a possible genetic interactions between DYF-5 and DYF-18.

**Figure 6.**
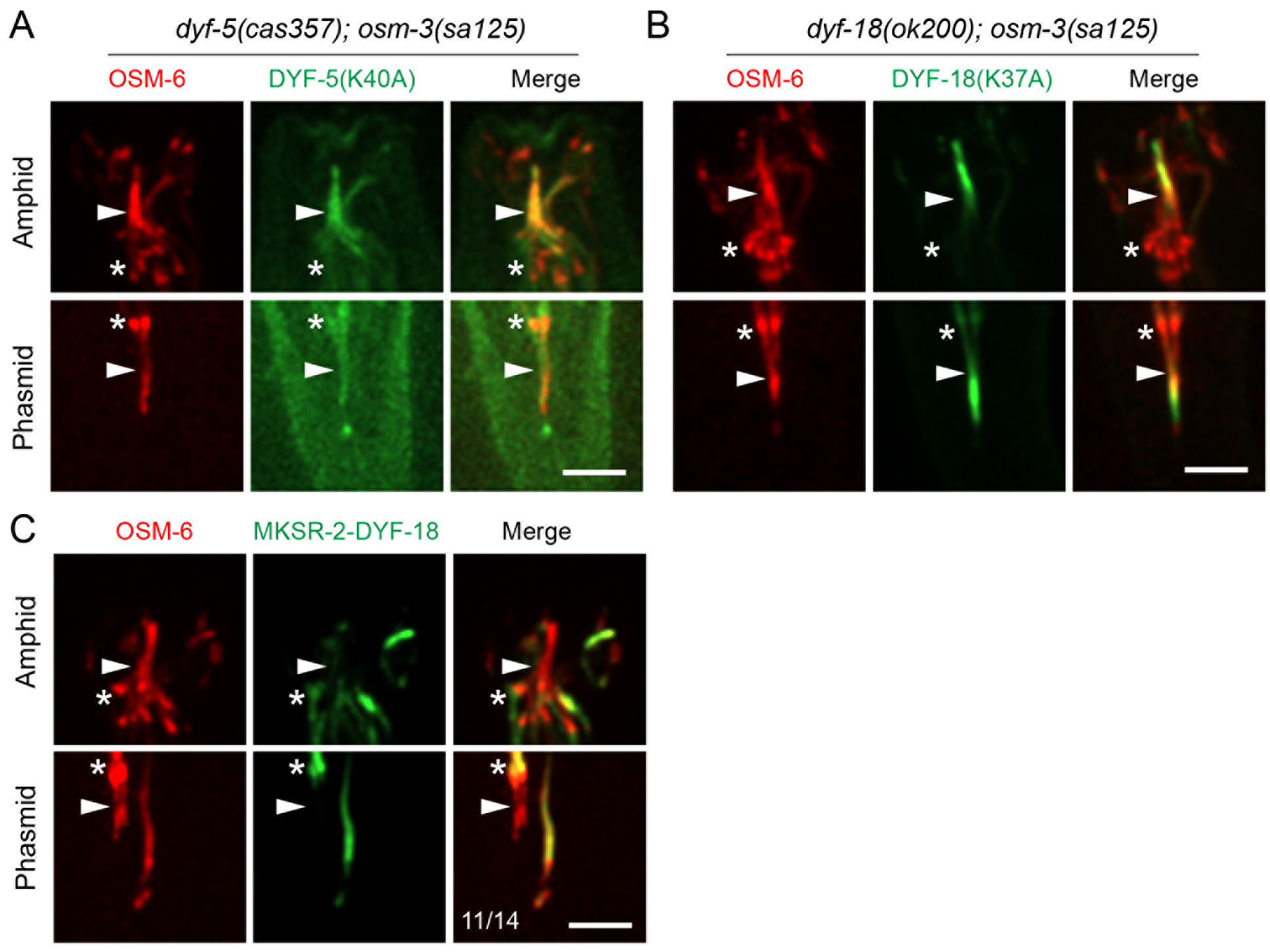
Kinase activity and ciliary localization are required for DYF-5 and DYF-18 function. (A-B) Kinase-dead DYF-5(K40A) or DYF-18(K37A) does not rescue the suppression of *osm-3(sa125)*. DYF-5a(K40A)::GFP, DYF-18(K37A) and OSM-6::mCherry are expressed under the control of the *Pdyf-1* promoter. Star, ciliary base; arrowhead, junction between the middle and distal segments; scale bar, 5 μm (the same in Figure 6C). (C) Localization of DYF-18 to ciliary base and transition zone causes the alignment defects in phasmid cilia (11 out of 14). Transition zone protein MKSR-2 was fused with DYF-18 and GFP. This chimeric construct was expressed under the control of the *Pdyf-1* promoter.

## Discussion

This study shows that the *C. elegans* homologs of the mammalian RCK kinases (MAK/ICK/MOK) and CCRK kinase, DYF-5 and DYF-18 are IFT cargo molecules and are enriched at the distal segments of sensory cilia. DYF-5 and DYF-18 play critical roles in coordinating the motility of kinsin-2 motor proteins and are involved in the recycling of IFT components.

### The interplay between DYF-5 and DYF-18

The partial inactivation of DYF-5 in *dyf-5* weak alleles displays similar ciliary phenotypes to those in the null allele of *dyf-18*; however, *dyf-5* null mutation causes strong ciliary phenotypes (Figure 2E & Figure 4G). These observations suggest that DYF-5 is essential for orchestrating anterograde and retrograde IFT whereas DYF-18 may act upstream of DYF-5 to modulate its activity. In supporting this view, genetic analyses in *Chlamydomonas* suggest that LF2/DYF-18 is epistatic to LF4/DYF-5 during flagellar regeneration (Asleson and Lefebvre, 1998); mammalian CCRK/DYF-18 directly phosphorylates the DYF-5 homologs ICK and MAK (Fu et al., 2006; Wang and Kung, 2012). The inhibition of glioblastoma cell proliferation by CCRK depletion is also found to depend on the ciliary functions of ICK and MAK (Yang et al., 2013). The different effects of *dyf-5* and *dyf-18* on ciliogenesis may reflect the intrinsic dual regulatory properties of DYF-5. DYF-5 belongs to the MAP kinase superfamily which contains a conserved TxY motif in the activation loop. Autophosphorylation of the tyrosine residue confers the basal kinase activity, and its full activity requires additional phosphorylation on the threonine residue by other kinases (Cobb and Goldsmith, 1995). This scenario is applicable to CCRK and ICK/MAK as evidenced by *in vitro* studies (Fu et al., 2006; Fu et al., 2005; Wang and Kung, 2012). Loss of *dyf-18* may represent the absence of full activation of DYF-5; however, the basal activity of DYF-5 may partially fulfil it functions in IFT and ciliogenesis as the *dyf-5* weak alleles. The enhanced ciliary defects in *dyf-18; dyf-5(weak allele)* double mutant may represent the additive effects on the loss of DYF-5 activity, which is evidenced by almost identical phenotypes in *dyf-18; dyf-5(weak allele)* double mutant and *dyf-5* null (Figure 4E-4G). Our imaging data further show that DYF-5 and DYF-18 have similar localization patterns, supporting the potentiality of their interaction *in vivo* (Figure 5).

We suggest that the ciliary function of DYF-5 requires precise control of its activity. Overexpression of *dyf-5* or its homologs in *L. Mexicana* and mammalian cells causes shorter cilia, whereas loss of its function induces longer cilia (Bengs et al., 2005; Broekhuis et al., 2014; Burghoorn et al., 2007; Moon et al., 2014). Loss or overexpression of CCRK causes similar phenotypes (Yang et al., 2013) and MAK and CCRK are found overexpressed in cancer cells where they promote proliferation by inhibiting ciliogenesis (Ng et al., 2007; Wang and Kung, 2012). CCRK is akin to the cyclin-dependent kinases (CDKs) in sequence and most similar to the CDK-activating kinase CDK7 (Wohlbold et al., 2006), implying that a CDK-like signaling cascade might function upstream of CCRK. Interestingly, the threonine in the activation loop of CCRK is replaced with an aspartate in DYF-18 (Figure S1B), which may explain why our overexpression of DYF-18 did not affect ciliogenesis in *C. elegans*. Besides CCRK and ICK/MAK, other cell cycle-associated kinases such as NIMA-related kinases CNK2 and CNK4 and CDK-like kinase CDKL5 are also found to regulate cilium length (Bradley and Quarmby, 2005; Hilton et al., 2013; Meng and Pan, 2016; Tam et al., 2013). We speculate that a cell cycle-related signaling network is diverged to inhibit ciliogenesis, although the relationship between these kinases remains to be addressed. Importantly, our suppressor screen also identified a NIMA kinase-like gene *nekl-4* which encodes the mammalian NEK10 homolog as a suppressor of *osm-3(sa125)* (Figure S3). NELK-4 localizes to the ciliary compartment and its loss of function only shows mild suppression phenotypes.

### DYF-5 and DYF-18 regulate the motility of kinesin-2 family proteins

How could DYF-5 affect the kinesin-II localization and recycling of IFT components? One possibility is that DYF-5 undocks kinesin-II by inactivating kinesin-II. The failure of kinesin-II undocking leads to increased IFT import to the distal segments, perturbing the balanced bidirectional IFT. However, OSM-3-driven anterograde IFT is also impaired in the absence of *dyf-5* (Burghoorn et al., 2007) and we cannot exclude the possibility that *dyf-5* is involved in retrograde IFT either. For example, DYF-5 may regulate the docking of IFT particles onto dynein-2, thus affecting dynein-2 motility. Considering that the activity of DYF-5 appears dosage dependent, the IFT and enrichment of DYF-5 at the ciliary distal segments suggest a cargo-dependent feedback that controls kinesin-II activity and cilium length (Figure 5). Interestingly, the phosphorylation and activation status of a kinase also correlates with flagella length in *Chlamydomonas* (Cao et al., 2013; Luo et al., 2011).

It is reasonable to speculate that DYF-5 inhibits kinesin-II activity via the direct phosphorylation of its subunits. To test this possibility, we performed GFP-based affinity purification of kinesin-II and mass spectrometry to identify phosphorylation sites. We determined two modified residues, Ser102 on KAP-1 and Ser735 on KLP-11 (Figure S4). However, neither the Ser-to-Ala nor Ser-to-Asp mutations could cause kinesin-II to localize at the distal segments. In addition, phosphorylation on these two residues are detectable in *dyf-5* null mutant. These data suggest that the phosphorylation of kinesin-II by DYF-5 regulation might be too low for detection; alternatively, DYF-5 may indirectly regulate kinesin-II activity by phosphorylating other IFT components. In zebrafish photoreceptors, IFT-B component IFT57 is required for the ATP-dependent dissociation of kinesin-II from the IFT particles (Krock and Perkins, 2008). Although the CDPK1-dependent phosphorylation and inactivation of kinesin-II was revealed, to our knowledge, no direct interactions between the *Chlamydomonas* LF4 and kinesin-II have been reported.

Our results suggest DYF-5 and DYF-18 regulate the handover between kinesin-II and OSM-3. Our GFP affinity purification and mass spectrometry analyses of GFP-tagged OSM-3 did not uncover any phosphorylation on OSM-3, raising the possibility of action of DYF-5 or DYF-18 on IFT particles to orchestrate these two motors. The *C. elegans* BBSome components could be potential candidates as the loss of *bbs-7* or *bbs-8* separates IFT-A and-B subcomplexes and BBSome modulates the competition between kinesin-II and OSM-3 (Ou et al., 2005; Pan et al., 2006). BBSome also interacts with IFT-A components to regulating turnaround of IFT particles (Wei et al., 2012). Other possible candidates include the OSM-3 activators, such as DYF-13, DYF-1,DYF-6, IFT-74 and IFT-81, among which IFT-74/IFT81 bind to OSM-3 (Hao et al., 2011). The handover between kinesin-II and OSM-3 can be coupled with the inactivation of kinesin-II and activation of OSM-3 through cargo undocking and docking, respectively. The bipartite axonemal structure might be also involved in regulating kinesin-2 motors, as *C. elegans* kinesin-II only moves on the doublet microtubules and returns at the tip of middle segments (Snow et al., 2004). Since *dyf-5* mutants contains singlet microtubules in the distal segments, the ectopic localization of kinesin-II might not be caused by altered axonemal microtubule structures (Burghoorn et al., 2007). However, phosphorylation of axonemal tubulins might be involved, as the motility and processitivity of kinesin-2 can be affected by posttranslationally modified tubulins *in vitro* (Sirajuddin et al., 2014).

## Materials and Methods

### Strain culture and genetics

All strains were raised on the nematode growth medium (NGM) plates seeded with *E. coli* OP50 at 20 °C. The genotypes of strains used in this study were listed in Table S1. To isolate *osm-3(sa125)* suppressors, *mnIs17[OSM-6::GFP]; osm-3(sa125)* mutant animals were synchronized at late L4 stage, collected in 4 mL M9 buffer, and incubated with 50 mM ethyl methanesulfonate (EMS) for 4 hours at room temperature with constant rotation. Animals were then washed with M9 for 3 times and recovered. 30 hours later, gravid animals were bleached. F1 eggs were divided and raised on ~100 9-cm NGM plates, each containing ~50 F1 eggs. Adult F2 animals on each plate were collected and subjected to dye filling (see Dye filling assay). Dye positive mutant animals were singly cultured and progenies were double-checked by dye filling assay and examination of ciliary phenotypes under microscopy. To map the mutations, a putative null allele *cas370* of *osm-3* (8 bp deletion at exon 2) was introduced into the CB4856 strain by CRISPR-Cas9-mediated knockout. Suppressor mutants were crossed with this strain to perform snip-mapping. Mutations were identified by whole genome and candidate sequencing. Isolated mutants were outcrossed for 3 times. Adult hermaphrodite worms were used in the dye filling assays and live imaging experiments. Worms at mixed stages were collected to perform GFP immunoprecipitation and mass spectrometry. All animal experiments were performed in accordance with the governmental and institutional guidelines.

### Molecular biology

The OSM-3(ΔH2) cDNA was a gift from Ronald Vale lab. OSM-3(G444E) coding sequence was amplified from *osm-3(sa125)* mutant genomic DNA. Constructs for overexpression of OSM-3(ΔH2), OSM-3(G444E), OSM-6, DYF-5a(K40A), DYF-18(K37A), TBB-4, NEKL-4, and kinesin-II subunits were generated by inserting their genomic coding sequences (OSM-3(G444E), OSM-6, DYF-5a(K40A), DYF-18(K37A), TBB-4) or cDNA (OSM-3(ΔH2), NEKL-4, KAP-1a and KLP-11a) into the pDONR vectors that contain the ~400 bp *dyf-1* promoter and GFP/mCherry*::unc-54* 3’UTR using In-Fusion Advantage PCR cloning kit (Clontech, cat. no. 639621). DYF-18::GFP or TBA-5::GFP fusion constructs were generated by SOEing PCR of the genomic sequences that contain their promoters and coding regions with the GFP::unc-54 3’UTR, respectively. ~900 bp promoter of *dyf-18* and ~5 kb promoter of *tba-5* were used. MKSR-2::DYF-18 chimeric expression vector was obtained by inserting the MKSR-2 genomic coding sequence into the *Pdyf-1::dyf-18::GFP* plasmid. Point mutations were introduced by transforming the linearized the vectors with 15 bp overlapping ends that bear necessary mutations into DH5α bacteria. For heat-shock induced expression, the *dyf-1* promoter was replaced with the ~400 bp *hsp-16.2* promoter. Primers used in this study were listed in Table S2.

### Knockout and knock-in

CRISPR-Cas9 targets were inserted to the pDD162 vector (Addgene #47549) in the same way as introducing point mutations. Target sequences for *dyf-18*, *kap-1* and *osm-3* knockout were AGGAAGTACCGTATACGTT(AGG), ATGCACACCCGTCAGATC(AGG), and ATGGCACTGTGTTTGCCTA(TGG) where the following protospacer adjacent motifs (PAMs) were shown in brackets. Two targets GAAGAAAGGCATGCGCCAAA(AGG) and CAATTGCAAACACTAAGAATAT(AGG) were used to generate DYF-5a::GFP knock-in strain. The homology arms are 850 bp (5’) and 1200 bp (3’) in length and were cloned into the pPD95.77 vector. GFP sequence was inserted prior to the stop codon. The last 11 amino acids of DYF-5a were removed to prevent the homology templates from Cas9 cutting. CRISPR-Cas9 constructs at the concentration of ~50 ng/μl were microinjected into the germ line of young adult hermaphrodites (*mnIs17[OSM-6::GFP]; osm-3(sa125)* mutant strain for *dyf-18* and *kap-1* knockout, CB4856 for *osm-3* knockout, and wild type N2 for *dyf-5* knock-in) together with *rol-6(su1006)* and *Podr-1::dsRed* markers. Roller and red fluorescence positive F1s were singly cultured. Successful edited animals were screened by PCR amplification and sequencing of mixed F2 progenies. Homozygous F2 animals without extrachromosomal transgenic markers were screened and verified by sequencing and outcrossed for 3 times. The *osm-3* mutant allele (in CB4856) *cas370* is an 8-bp deletion “GTTTGCCT” at exon 2, the *kap-1* allele *cas399* is a 14-bp deletion “CGTCAGATCAGGCA” at exon 1, and the *dyf-18* allele *cas444* is a compound mutation that bears a 3-bp deletion and a 58-bp insertion “<u>TACCGT</u>ATA/TAGGGGATATTATTAAAGACAAAACACGACCTGATGTCAATT GTTACCGAGGAAGTAC<u>CGTTAG</u>” (flanking sequences are underlined) at exon 3. All three mutations introduce frame shifts and early premature stop codons, thus are putative null alleles.

### Transgenesis

Transgenic lines of OSM-4(G444E)::GFP, OSM-3(ΔH2)::GFP, TBA-5::GFP, TBB-4: mCherry, DYF-18::GFP, DYF-5(K40A)::GFP, DYF-18(K37A)::GFP, MKSR-2::DYF-18, NEKL-4::GFP, and kinesin-II::GFP/mCherry were obtained by injecting these constructs with appropriate selection markers (*rol-6(su1006)* or P*dyf-1::osm-6::mCherry*) into the germ line of wild type N2 or mutant young adult animals. Marker or fluorescence positive F1s were singly cultured and maintained if F2s inherited the transgenes. Concentration of each DNA construct for microinjection was ~50 ng/μl. Similar expression patterns were observed in at least two or three independent lines of each transgenic strain.

### Dye filling assay

Adult animals were collected into ~200 μl M9 buffer, washed for 2 times, and mixed with equal-volume fluorescent dye (DiI 1,1’-dioctadecyl-3,3,3’,3’,-tetramethylindo-carbocyanine perchlorate, Sigma) at the concentration of 20 μg/ml. After incubation at room temperature in dark for 30-60 min, animals were transferred to seeded NGM plates to excrete excess intestinal dye. One hour later, animals were examined for dye filling in head and tail sensory neurons using compound microscope.

### Live imaging and analysis

Adult animals were immobilized with 0.1 mmol/L levamisole in M9 buffer on 3% agar pads in rounded dishes and maintained at room temperature for imaging of ciliary morphology and IFT. Our imaging system includes an Axio Observer Z1 microscope (Carl Zeiss) equipped with a 100×, 1.46 NA objective lens, an EMCCD camera (iXon+ DU-897D-C00-#BV-500; Andor Technology), and the 405 nm, 488 nm and 568 nm lines of a Sapphire CW CDRH USB Laser System (Coherent) with a spinning disk confocal scan head (CSU-X1 Spinning Disk Unit; Yokogawa Electric Corporation). Images were acquired by μManager (http://www.micro-manager.org). Still images of ciliary morphology were taken at an exposure time of 100-200 milliseconds and time-lapse images showing IFT were acquired at an exposure time of 200 milliseconds. Laser power was 20-30%. Kymograph extraction, image processing, and measurement of cilium length and IFT speeds were performed in ImageJ (http://rsbweb.nih.gov/ij/).

### Bioinformatics

Sequence alignment was performed using Clustal X2.1 (http://www.clustal.org/). Protein sequences were obtained from Wormbase (http://www.wormbase.org/) or National Center for Biotechnology Information (http://www.ncbi.nlm.nih.gov/). Conserved domains were identified by SMART (simple modular architecture research tool) (http://smart.embl-heidelberg.de/) or CDD (NCBI’s conserved domain database) online tools. Sequence IDs used include CAQ76489.2 (*Ce*DYF-5), P20794.2 (*Hs*MAK), NP_055735.1 (*Hs*ICK), BAA81688.1 (*Hs*MOK), AAO86687.1 (*Cr*LF4), CAA43985.1 (*Hs*CDK2), CAB07422.2 (*Ce*DYF-18), NP_001034892.1 (*Hs*CCRK), ABK34486.1 (*Cr*LF2), NP_001790.1(*Hs*CDK7), NP_001186326.1 (*Hs*NEK1), NP_002488.1(*Hs*NEK2), NP_002489.1 (*Hs*NEK3), NP_003148.2 (*Hs*NEK4), NP_954983.1 (*Hs*NEK5), CAG33372.1 (*Hs*NEK6), BAB85632.1 (*Hs*NEK7), AAP04006.1 (*Hs*NEK8), NP_149107.4 (*Hs*NEK9), NP_689747.3 (*Hs*NEK10), NP_079076.3 (*Hs*NEK11), NP_001293320.1 (*Ce*NEKL-1), NP_491914.1 (*Ce*NEKL-2), NP_510080.2 (*Ce*NEKL-3), and NP_498178.3 (*Ce*NEKL4). Full sequences were used to perform alignment except *Hs*NEK10 and *Ce*NEKL-4 whose kinase domains were used.

### Quantification and statistical analysis

Quantification was represented by the mean value ± standard deviation for each group. Two-tailed student’s *t* test analysis was performed to examine all significant differences between groups as indicated in the figure legends. Numbers of subjects (animals, cilia or kymographs) used for quantification were indicated in the figure legends. Significance was determined when the *p* value is lower than 0.05. The variance is not significantly different between groups that are being compared unless it is mentioned in the text.

## Author Contributions

G. Ou and P. Yi conceived this project. P. Yi and C. Xie performed the experiments. P. Yi collected and analyzed the data under the supervision of G. Ou. G. Ou and P. Yi wrote the manuscript.

## Acknowledgements

We thank Caenorhabditis Genetics Center (CGC) for providing some strains and the Protein Chemistry Facility at the Center for Biomedical Analysis of Tsinghua University for sample analysis. This work was supported by the National Basic Research Program of China (grant 2017YFA0503501), the National Natural Science Foundation of China (grants 31525015 and 31561130153) and the Newton Advanced Fellowship from the Royal Society.

The authors declare no competing financial interests.

---

**Figure S1. Sequence alignments of mutated residues and the activation loops of DYF-5 and DYF-18**.

(A) Sequence alignment of mutated residues in *C. elegans* (*Ce*) DYF-5 and DYF-18 and their homologs in human (*Hs*) and *Chlamydomonas* (*Cr*). Arrows indicate mutations isolated by EMS screening (also see Figure 2B & 2C).

(B) Sequence alignment of the activation loops of DYF-5 and DYF-18. Stars mark the threonine residues subject to phosphorylation. The conserved TxY motif in MAP kinase family is indicated.

**Figure S2. Mutations of *dyf-5* and *dyf-18* suppress the axonemal length defects in *osm-3(sa125)***.

(A) Ciliary phenotypes in *osm-3(sa125)* mutant. Cilia are marked with GFP labeled α-tubulin TBA-5 and mCherry labeled (β-tubulin TBB-4. TBA-5::GFP is expressed under its native promoter. TBB-4::mCherry is expressed under the control of the *Pdyf-1* promoter. Star, ciliary base; arrowhead, junction between the middle and distal segments; scale bar, 5 μm (the same in Figure S2B & S2C).

(B) Ciliary phenotypes in *osm-3(sa125); dyf-5(cas357)* double mutant.

(C) Ciliary phenotypes in *osm-3(sa125); dyf-18(cas390)* double mutant.

**Figure S3. Mutations of *nekl-4* suppress ciliary phenotypes in *osm-3(sa125)* mutant**.

(A) Genomic structure of *nekl-4*. Exons are marked with boxes. NEKL-4 belongs the NIMA-kinase superfamily and homologous to mammalian NEK10. Blue regions indicate the coding sequence of kinase domain. Scale bar, 500 bp.

(B) Sequence alignment of the human (*Hs*) and *C. elegans* (*Ce*) NIMA-kinase family members. Star indicates the *cas396* mutation which converts a hydrophobic alanine to a polar threonine residue.

(C) Ciliary phenotypes in *nekl-4(cas396); osm-3(sa125)* double mutant. Cilia are marked with GFP-tagged IFT52/OSM-6. Star, ciliary base; arrowhead, junction between the middle and distal segments; scale bar, 5 μm.

(D) Ciliary localization of NEKL-4. NEKL-4::GFP and OSM-6::mCherry are overexpressed under the control of the *Pdyf-1* promoter. Star, ciliary base; arrowhead, junction between the middle and distal segments; scale bar, 5 μm.

**Figure S4. Mutations of two phosphorylated residues in kinesin-II do not affect its localization**.

(A) Domain structures of the *C. elegans* kinesin-II accessory subunit isoform a (KAP-1a) and motor subunit isoform a (KLP-11a). KAP domain is in blue. The motor domain and coiled-coil domain of KLP-11 are in pink and green, respectively. The phosphorylated residues (Ser102 of KAP-1 and Ser735 of KLP-11) are indicated.

(B) Ser-to-Ala mutation does not alter KAP-1 or KLP-11 localization. KAP-1a(S102A)::mCherry and KLP-11a::GFP are expressed under the control of the *Pdyf-1* promoter. Star, ciliary base; arrowhead, junction between the middle and distal segments; scale bar, 5 μm.

(C)Simultaneous Ser-to-Ala or Ser-to-Asp mutations on KAP-1 and KLP-11 do not alter kinesin-II localization. KAP-1a(S102A)::mCherry and KLP-11a::GFP are expressed under the control of the *Pdyf-1* promoter. Star, ciliary base; arrowhead, junction between the middle and distal segments; scale bar, 5 μm.

**Table S1.**
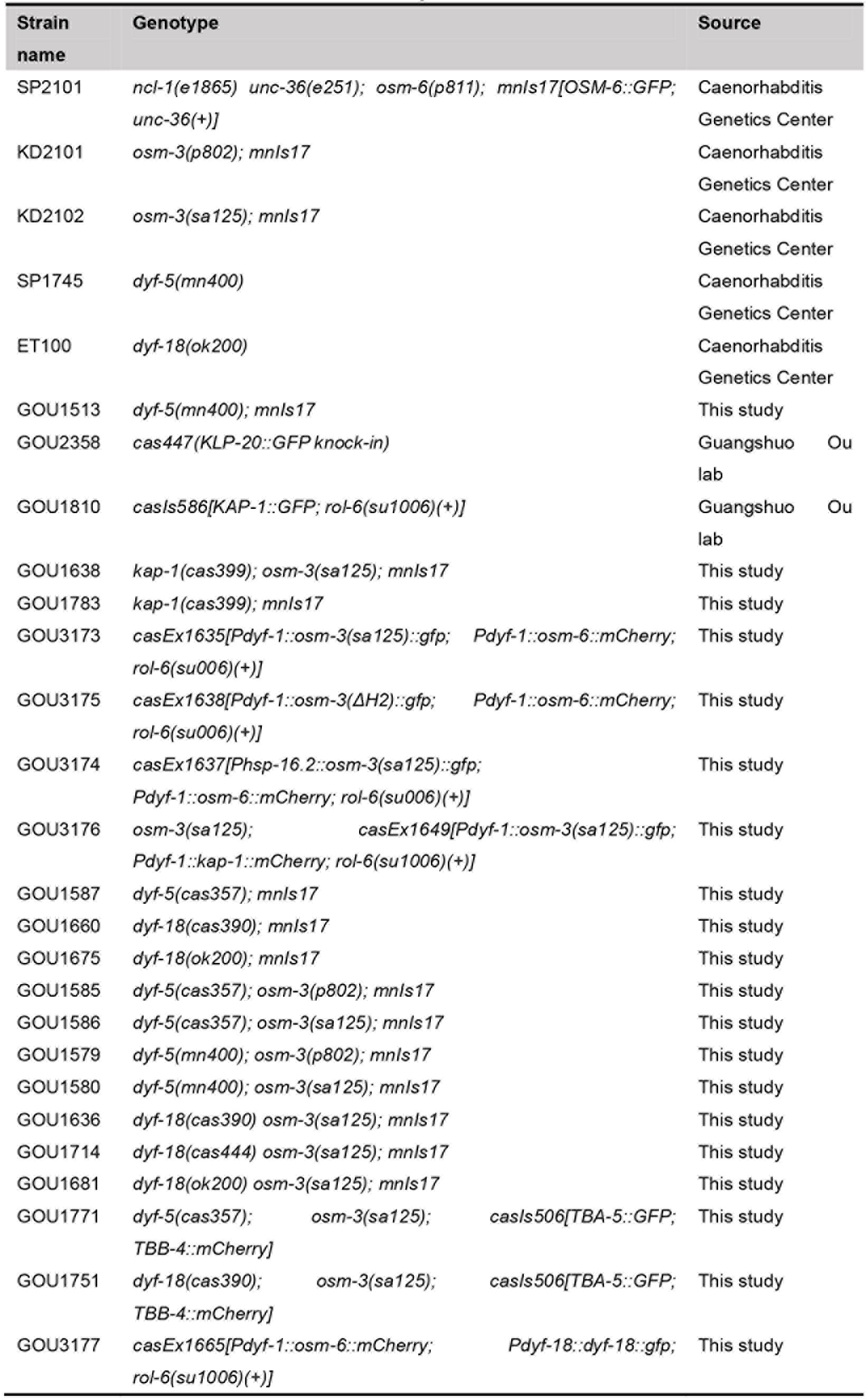
**Strains used in this study**.

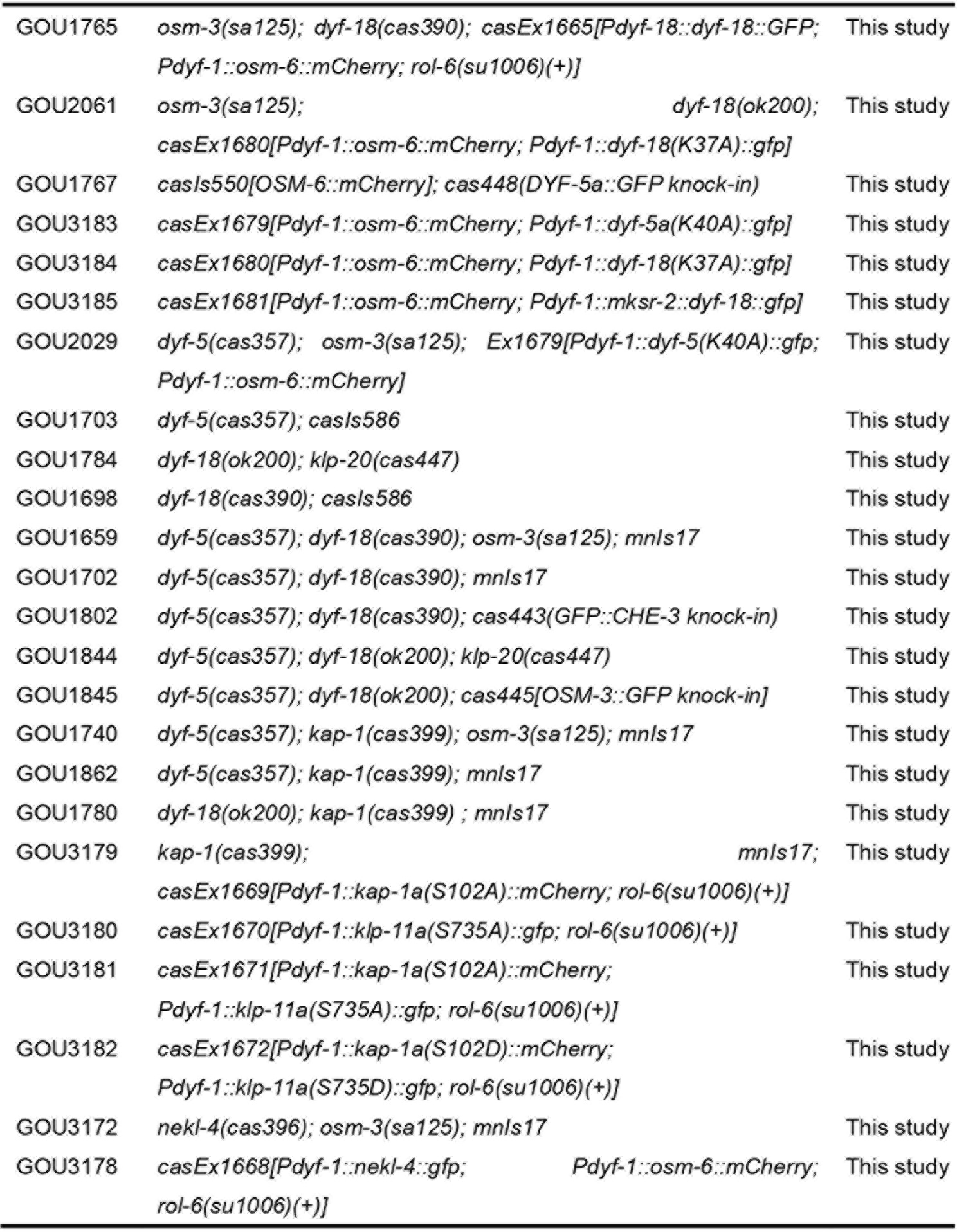

**Table S2.**
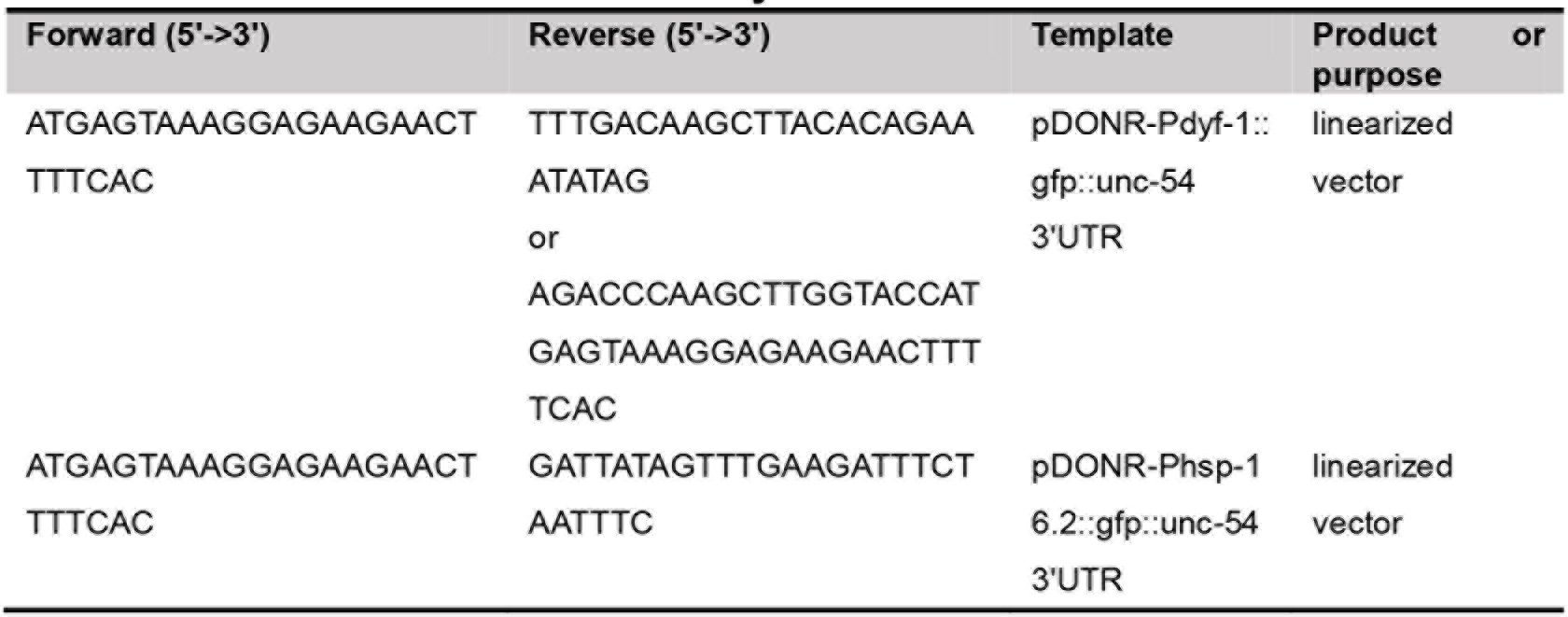
**Primers used in this study**.

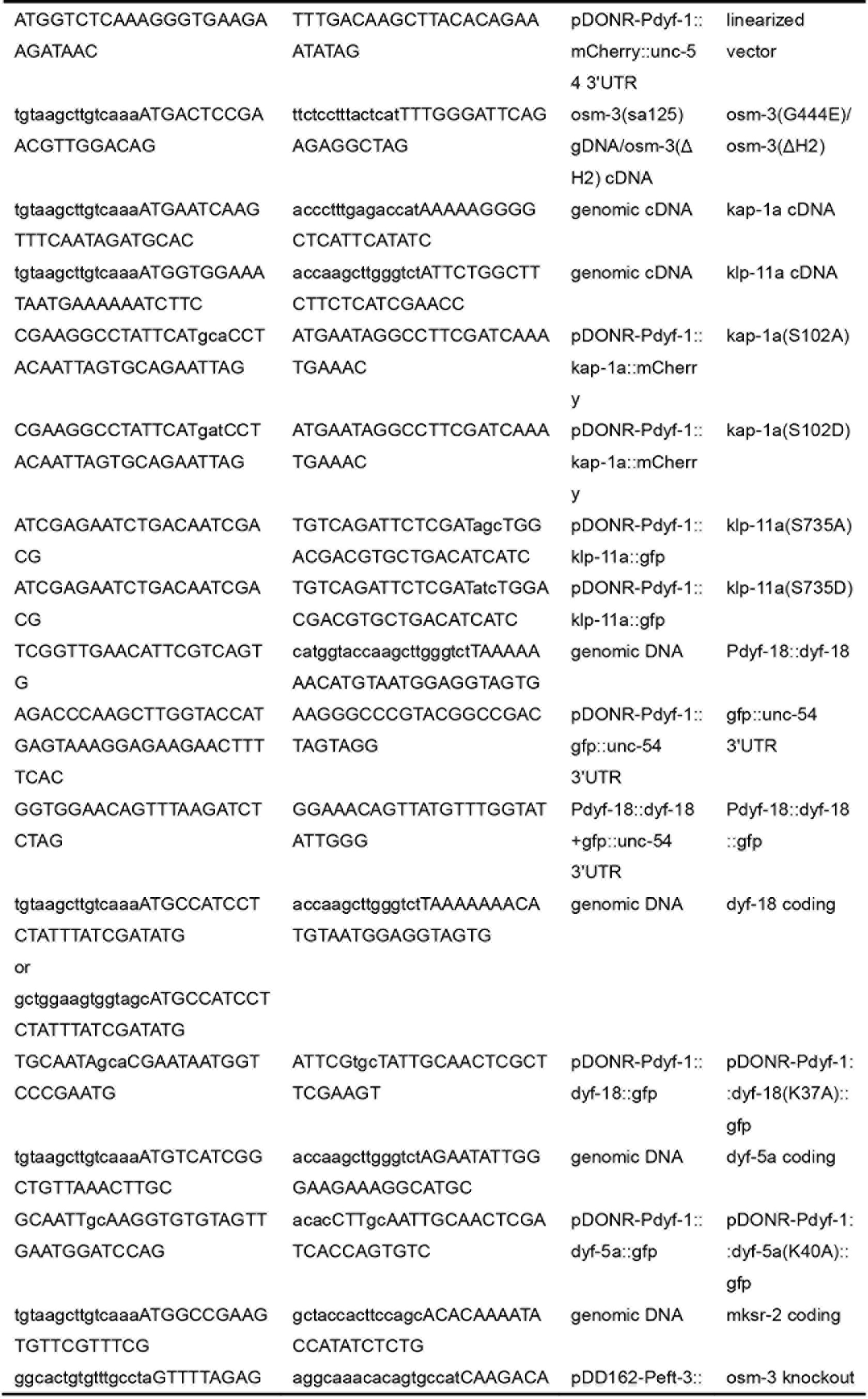

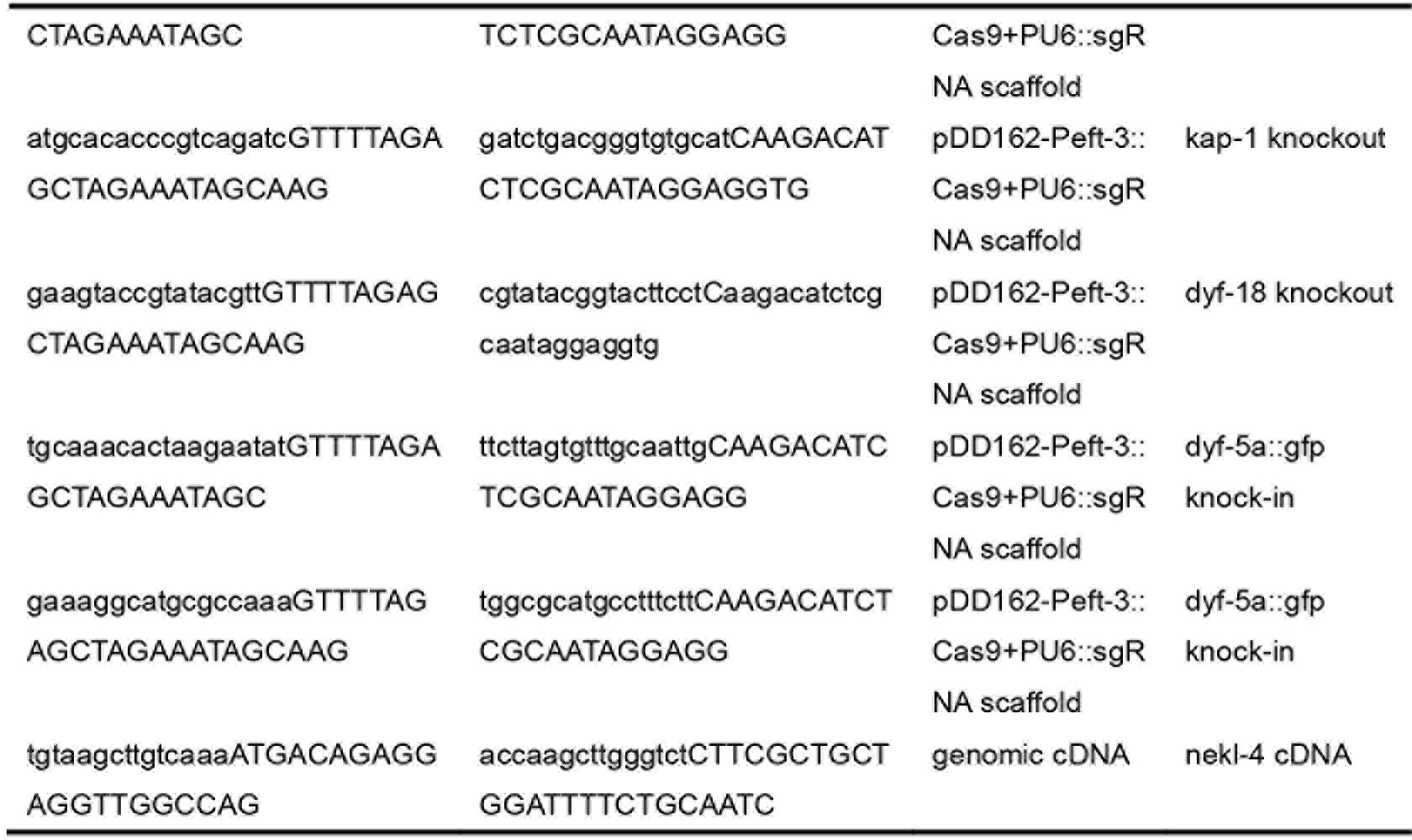

